# “Proliferation of DBLOX Peroxidase-Expressing Oenocytes Maintains Innate Immune Memory in Primed Mosquitoes”

**DOI:** 10.1101/2020.09.09.290312

**Authors:** Fabio M. Gomes, Miles D.W. Tyner, Ana Beatriz F. Barletta, Lampougin Yenkoidiok-Douti, Gaspar E. Canepa, Alvaro Molina-Cruz, Carolina Barillas-Mury

**Author notes:** These authors contributed equally to this work (Fabio M. Gomes, Miles D.W. Tyner).

## Abstract

Immune priming in *Anopheles gambiae* mosquitoes following infection with *Plasmodium* parasites is mediated by the systemic release of a hemocyte differentiation factor (HDF), a complex of lipoxin A_4_ bound to Evokin, a lipid carrier. HDF increases the proportion of circulating granulocytes and enhances mosquito cellular immunity. We found that Evokin is constitutively produced by hemocytes and fat-body cells, but expression increases in response to infection. Insects synthesize lipoxins, but lack lipoxygenases. Here, we show that the Double Peroxidase (DBLOX) enzyme, present in insects but not in vertebrates, is essential for HDF synthesis. DBLOX is highly expressed in oenocytes in the fat body tissue, and these cells proliferate in response to *Plasmodium* challenge. We provide direct evidence that modifications mediated by the histone acetyltransferase AgTip60 (AGAP01539) are essential for sustained oenocyte proliferation, HDF synthesis and immune priming. We propose that oenocytes function as a population of “memory” cells that continuously release lipoxin to orchestrate and maintain a broad, systemic and long-lasting state of enhanced immune surveillance.

## Main Text

There is growing evidence that a previous infection can “train” or “prime” the innate immune system, allowing it to respond more effectively to subsequent infections (*1–3*). This is an ancient response that has been documented in plants (*4*), insects (*5*) and humans (*6*). It is not clear whether individual immune cells maintain the training response, or if there are specific “memory keeper” cell populations that orchestrate a persistent re-training of effector immune cells. In *Anopheles gambiae*, the primary vector of malaria in Africa, *Plasmodium* infection induces a long-lasting priming response that enhances antiplasmodial immunity (*5*). *Plasmodium* midgut invasion allows direct contact between the gut microbiota and midgut epithelial cells, triggering a burst of prostaglandin E_2_ (PGE_2_) production by midgut cells. The transient systemic release of PGE_2_ establishes a long-lasting release of hemocyte differentiation factor (HDF) which, in turn, induces an increase in the proportion of circulating granulocytes, a hallmark of immune priming (*7, 8*). HDF is a complex of Evokin, a lipocalin lipid carrier protein, and the lipid mediator Lipoxin A_4_ (LXA_4_) (*7*). Thus, immune priming can be defined as the enhanced ability of mosquitoes to convert arachidonic acid into LXA_4_ that results in a permanent functional state of enhanced immune surveillance (*7*). An effective antiplasmodial response requires the coordinated activation of epithelial, cellular, and complement components of the mosquito immune system (*9*). PGE_2_ attracts hemocytes to the basal surface of the midgut (*8*), and primed hemocytes release more microvesicles, enhancing complement-mediated ookinete lysis (*9*).

In vertebrates, cyclooxygenases (COX) and lipoxygenases (LOX) catalyze the synthesis of prostaglandins and lipoxins, respectively (*10*). Although eicosanoids have been detected in mosquitoes and other insects (*8, 11, 12*), neither COX nor LOX enzymes are present in insects. We recently showed that two heme peroxidases, HPX7 and HPX8, are necessary for midgut PGE_2_ synthesis and are essential to establish immune priming in response to *Plasmodium* infection (*8*). However, the enzyme(s) mediating LXA_4_ synthesis in insects remain unknown. In this manuscript, we identify the Double Peroxidase (DBLOX) enzyme as essential for HDF synthesis, and show that DBLOX is highly expressed in oenocytes, a cell population involved in lipid processing. Furthermore, we discovered that oenocytes proliferate in primed mosquitoes. In vertebrates, monocyte trained immunity is mediated by epigenetic modifications (*13, 14*). Here, we identify a histone acetyltransferase (HAT) that is an essential mediator of oenocyte proliferation and mosquito immune priming.

mRNA expression of Evokin, an essential component of HDF (*7*), was significantly induced in the hemocyte-like immune-responsive *A. gambiae* Sua 5.1 cell line in response to bacterial challenge (Fig. 1A) (p=0.0037, Mann-Whitney test). Additionally, antibodies to recombinant Evokin recognized a single band of the expected size (21 kDa) in the fraction containing membrane and insoluble proteins, but not in the cytoplasm (Fig. 1B). Evokin mRNA expression also increased significantly in hemocytes (p=0.0006, unpaired t-test) and body wall (p= 0.0005, unpaired t-test) of primed mosquitoes 7 days post-infection (Fig 1C). In naïve blood-fed females, Evokin is highly expressed in sessile hemocytes associated with the body wall surface (Fig 1D). It is also present in oenocytes, with a dotted vesicle-like pattern (Fig 1E), while in fat body trophocytes a string pattern is observed, suggestive of Evokin clustering in specific membrane regions that may represent lipid rafts (Fig 1F). A similar expression pattern was observed in *Plasmodium* challenged females (Fig. S1).

**Fig 1.**
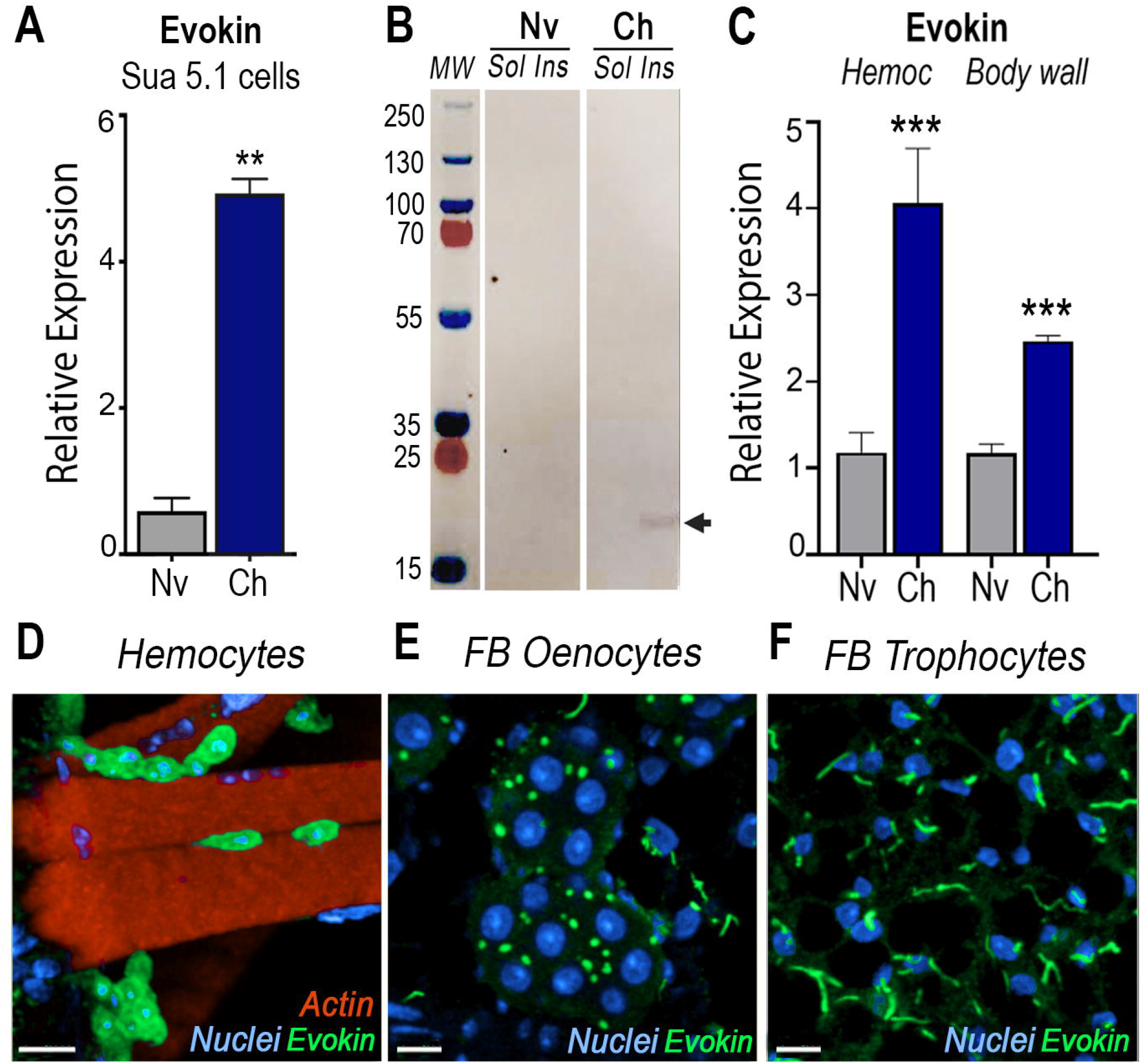
Effect of immune challenge of Sua 5.1 cells and mosquitoes on Evokin expression. Effect of bacterial challenge on Sua 5.1 cells on (**A**) Evokin mRNA levels measured by qRT-PCR and (**B**) Evokin protein expression (arrow) analyzed by Western Blot. (**C**) Effect of *P. berghei* infection on Evokin mRNA levels measured by qRT-PCR in mosquito hemocytes and body wall 7 days post-infection. Locatization of evokin in (**D**) sessile, body wall-associated hemocytes, (**E**) oenocytes, and (**F**) trophocytes analyzed by immunofluorescence staining of naïve mosquitoes. Mean ± SEM are plotted and groups were compared using Student’s t test (**p<0.01, ***p<0.001). Scale bars 7 μm (**D**), 10 μm (**E**), 10 μm (**F**). Nv, naïve; Ch, challenged; Sol, soluble fraction; Ins, insoluble fraction; Hem, hemocytes.

Given the critical role of HPX7 and HPX8 in PGE_2_ synthesis, we investigated if other HPX enzymes could mediate the conversion of arachidonic acid to LXA_4_. Changes in mRNA expression of 15 HPX enzymes, DBLOX and Dual Oxidase (DUOX) in mosquitoes primed by *P. berghei* infections (Fig. 2A) or by systemic injection of PGE_2_ (Fig. 2B) were evaluated (Table S1). Both treatments resulted in a prolonged and significant increase in DBLOX expression in the body wall (p=0.0056 and p=0.0059, respectively, Mann-Whitney test) while HPX2 was only induced following PGE_2_ injection (p=0.0058, Mann-Whitney test) (Fig. 2B). HPX2 silencing did not affect priming, as the characteristic increase in granulocytes was observed in response to *Plasmodium* infection (p=0.0001, Mann Whitney test) (Fig. 2C). In contrast, DBLOX silencing completely abolished the priming response to infection (Fig. 2D) and to PGE_2_ injection (Fig. S2). Furthermore, hemolymph of mosquitoes in which DBLOX was silenced no longer had HDF activity when transferred to naïve mosquitoes (Fig. 2E).

**Fig 2.**
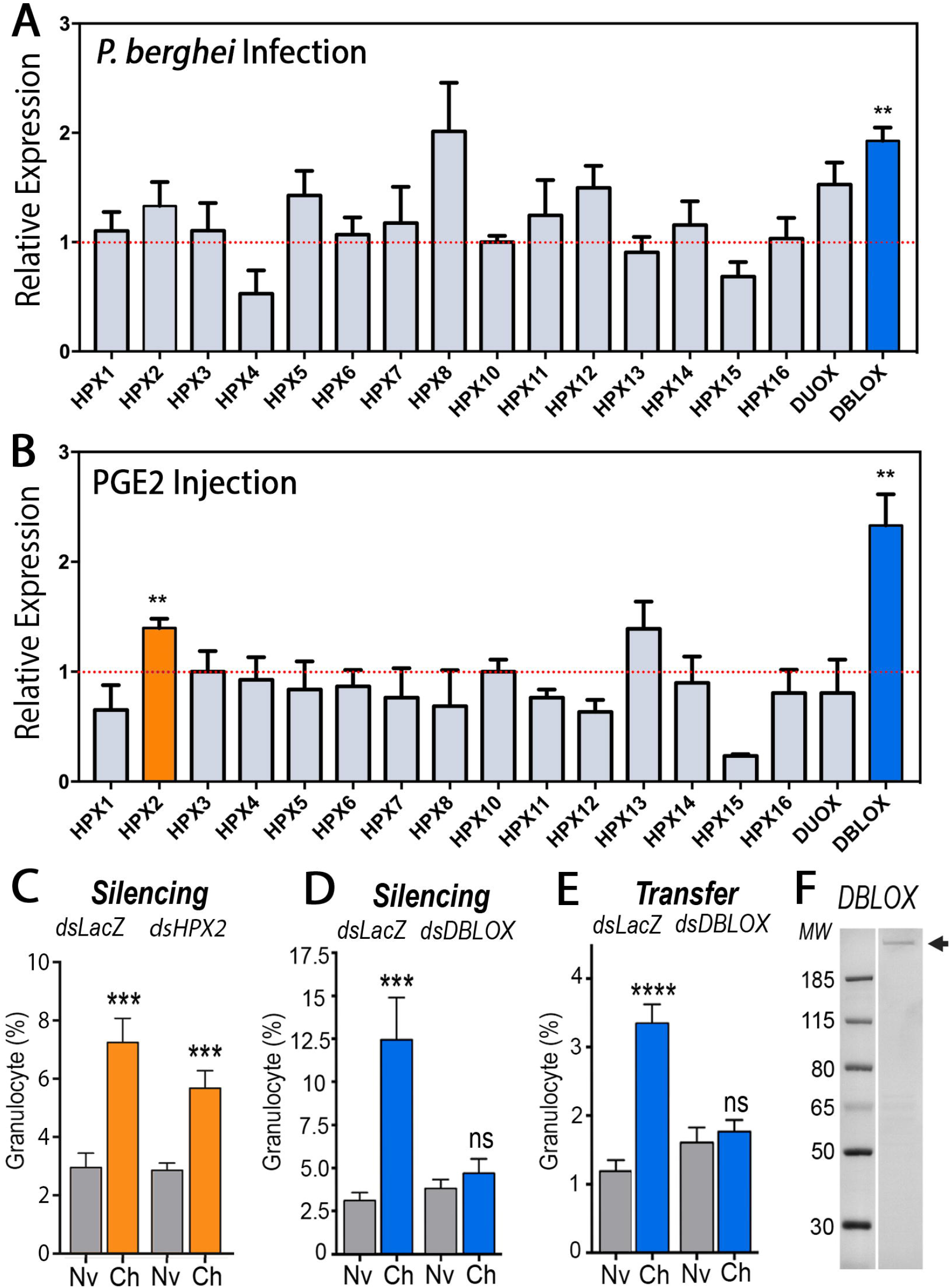
Effect of *Plasmodium* infection and PGE_2_ injection on heme peroxidases (HPXs) expression and role of HPXs on immune priming. (**A**) HPX mRNA expression 7 days after *P. berghei* infection and (**B**) 6 days after PGE_2_ injection. Effect of silencing (**C**) HPX2 and (**D**) DBLOX on *Plasmodium*-induced granulocyte priming. (**E**) Effect of silencing DBLOX on *Plasmodium*-induced HDF activity. (**F**) DBLOX protein expression (arrow) from hemocyte-like Sua 5.1 cells in response to bacterial challenge analyzed by Western Blot. Mean ± SEM are plotted and groups were compared using Student’s t test (**p<0.01, ***p<0.001, ****p<0.0001). Nv, naïve; Ch, challenged.

Antibodies to recombinant DBLOX detected a single band under native conditions (Fig 2F) of the expected size (159 kDa) (Fig S3). DBLOX is highly expressed in oenocytes that form clusters on the fat body surface (Figs. 3A-B). The DBLOX staining is cytoplasmic, with a punctate pattern and is also present in lower abundance in the perinuclear region of fat body trophocytes (Figs. 3A-B). Oenocytes proliferated in primed mosquitoes 4 days post-infection with a significant increase in both the number of cell clusters (p = 0.0031, Mann Whitney test,) and number of cells per body wall segment (p = 0.0001, Mann Whitney test) (Figs. 3C-E). This proliferation is long-lasting, as the increased number of clusters and cells per segment were still present 7 days post-feeding (Fig. S4). Given the essential role of DBLOX in HDF synthesis and immune priming, the high expression of DBLOX in oenocytes, and their sustained proliferation in response to priming, we infer that the documented enhanced ability of primed mosquitoes to synthesize LXA_4_ from arachidonic acid (*7*) is achieved by increasing the number of oenocytes, the cells in the mosquito that are a major site of LXA_4_ synthesis.

**Fig 3.**
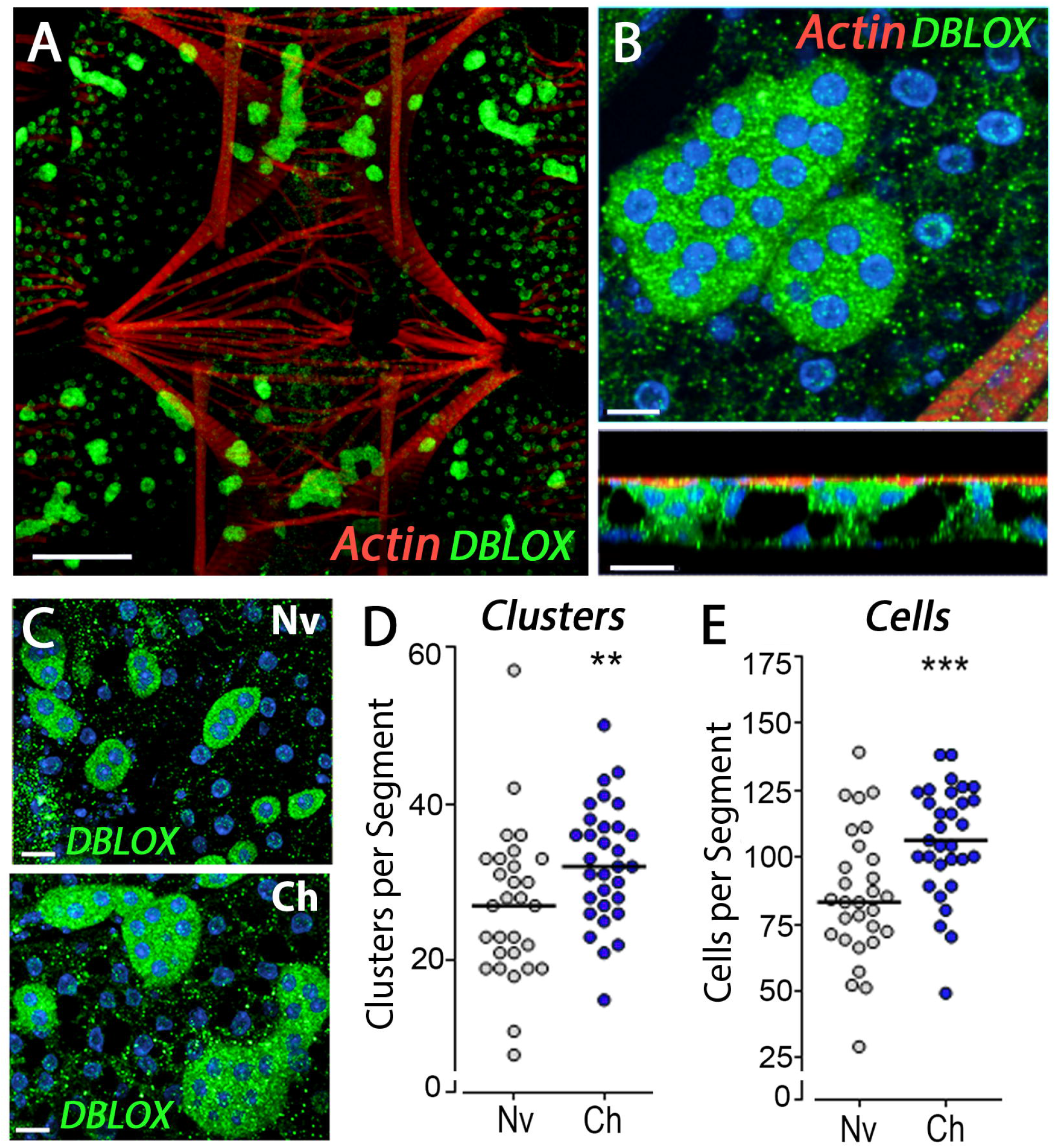
Tissue localization of DBLOX expression and effect of *Plasmodium* infection on mosquito fat body eonocytes. DBLOX expression by immunofluorescence staining in (**A**) the ventral and lateral body wall, and (**B**) oenocyte clusters at higher magnification (upper panel) and side view (lower panel). Effect of *Plasmodium* infection on (**C**) DBLOX-expressing oenocytes revealed by immunofluorescence staining, (**D**) the number of oenocyte clusters per body wall segment, and (**E**) the number of oenocytes per body wall segment. Lines indicate medians that were compared using the Mann-Whitney U test. (**p<0.01, ***p<0.001). Scale bars 100 μm (**A**), 10 μm (front view) and 20 μm (side view) (**B**), 15 μm (C). Nv, naïve; Ch, challenged.

DBLOX and Evokin mRNA levels remain persistently high in mosquitoes primed through *P. berghei* challenge or by PGE_2_ injection. We explore whether the establishment and persistence of immune priming is mediated by histone acetyltransferases (HAT), enzymes known to catalyze epigenetic chromatin modifications associated with long-lasting changes in transcriptional regulation (*15, 16*). We evaluated the effect of silencing each of the ten HATs present in the *A. gambiae* genome (AgHATs) on immune priming following *P. berghei* infection (Fig. 4A). Silencing AGAP000029 resulted in high mortality after blood-feeding and, thus, it could not be further evaluated. Of the other nine AgHATs, only silencing of the homolog of *Drosophila* Tip60 (AGAP001539 - AgTip60) abolished immune priming (Fig. 4A). We confirmed that dsLacZ injection had no effect on priming, while the proportion of granulocytes no longer increased when AgTip60 was silenced (Fig. 4B). Furthermore, hemolymph of primed AgTip60-silenced females no longer had HDF activity when transferred to naïve mosquitoes (Fig. 4C). In agreement with our proposed model, the loss of HDF activity following AgTip60 silencing was also associated with a lack of proliferation of DBLOX-expressing oenocytes, as the number of clusters and cells per segment no longer increased following a challenge with *P. berghei* (Fig. 4D-F).

**Fig 4.**
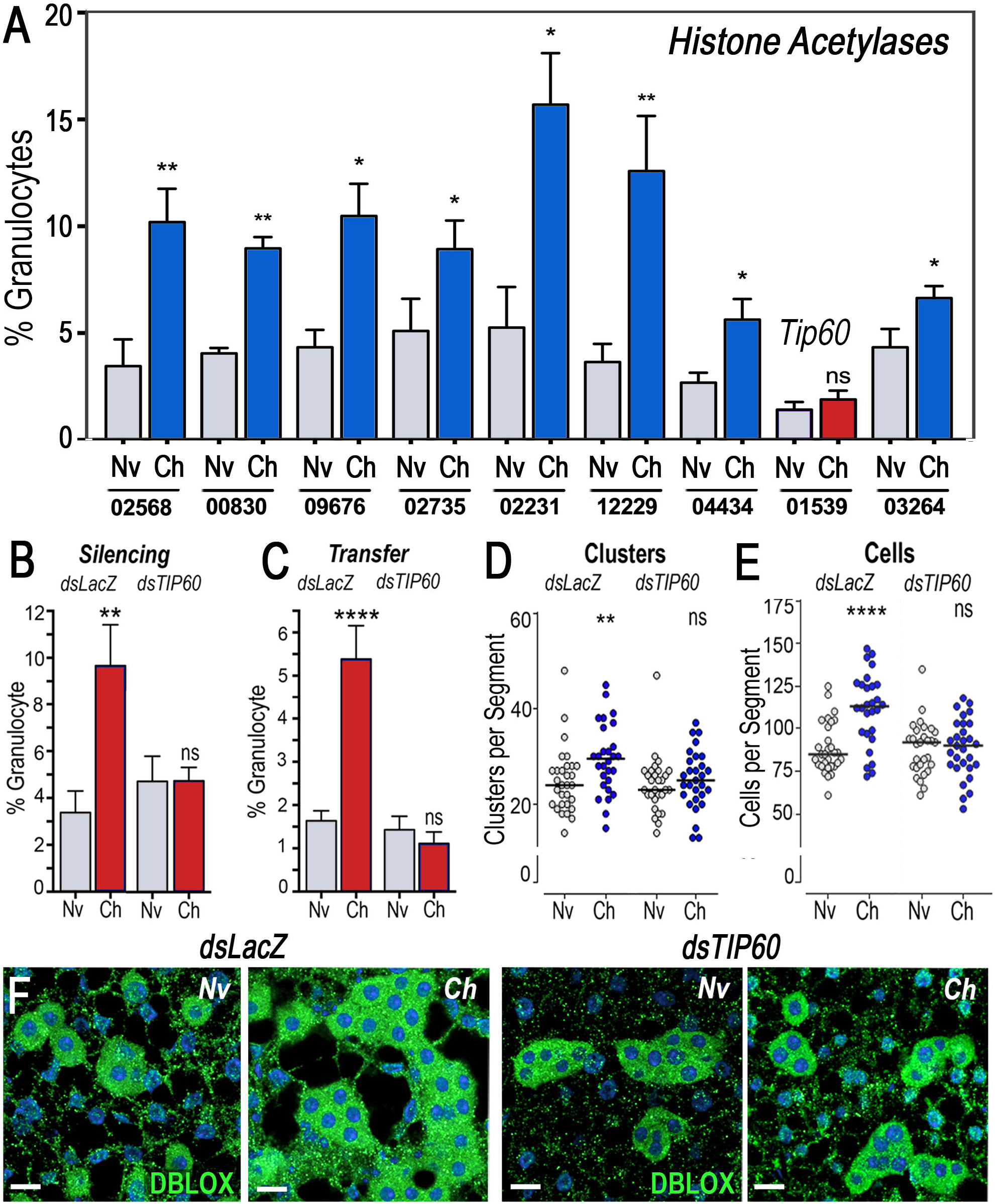
Effect of histone acetyltransferases (HATs) silencing on priming and proliferation of DBLOX-expressing oenocytes. (**A**) Effect of silencing each *A. gambiae* histone HATs on granulocyte priming. Numbers at the bottom indicate the AGAP reference number of each silenced gene. Effect of silencing AgTip60 (AGAP01539) on (**B**) *Plasmodium*-induced granulocyte priming, (**C**) HDF activity, (**D**) number of oenocyte clusters and (**E**) number of oenocytes. (**F**) Effect of silencing AgTip60 (TIP60) on immunofluorescence staining of DBLOX in body wall of naïve and challenged mosquitoes. Scale bars 15 μm. Mean ± SEM are plotted and groups were compared using Student’s t test (*p<0.05, **p<0.01, ****p<0.0001). Nv, naïve; Ch, challenged.

DBLOX is a unique enzyme with two duplicated heme peroxidase domains that is present in insects, but not in vertebrates. The first domain has the predicted substrate binding sites but lacks the functional residues present in catalytically active enzymes and has two integrin-binding motifs, typical of peroxinectins (*17*). In contrast, the second domain has all the features of a functional heme peroxidase and one integrin-binding motif (*17*). DBLOX is essential for LXA_4_ synthesis, but the biochemical mechanism of lipoxygenase-independent lipoxin synthesis in insects, and the potential involvement of other enzymes besides DBLOX remain to be explored. We speculate that the systemic burst of PGE_2_ when ookinete invasion allows direct contact between the microbiota and gut epithelial cells, or when mosquitoes are injected with PGE_2_, triggers modification mediated by AgTip60 that establish and maintain proliferation of oenocytes. These cells express high levels of DBLOX and their proliferation is essential for HDF synthesis and to maintain the priming response. The function of oenocytes is poorly understood, but they are known to be involved in biosynthesis of cuticular hydrocarbon and pheromones (*18*). Taken together, our findings suggest that oenocytes are a major site of lipoxin synthesis.

Immune training of human monocytes involves transcriptional and metabolic reprogramming mediated by epigenetic modifications that allow challenged monocytes to maintain a prolonged state of enhanced immune function (*13, 19*). We propose that oenocytes function as “memory keeper” cells that persistently release LXA_4_, which constantly re-trains mosquito hemocytes. Recently, trained immunity in humans induced by vaccination with the anti-tuberculosis bacillus Calmette-Guérin (BCG) vaccine has received much attention in light of the COVID19 pandemic (*20*). There is strong epidemiological evidence that BCG vaccination in infants has a broad protective effect and reduces the overall mortality from respiratory infections not related to tuberculosis (*21–23*), and there are ongoing clinical studies to establish if the BCG vaccine also protects from severe COVID19 (*24, 25*). Furthermore, there is growing epidemiological evidence of reduced COVID19 mortality (*26*) and improved outcomes (*27*) in countries with strong BCG vaccination programs. However, little is known about the mechanism to establish and maintain trained immunity. In *A. gambiae*, prostaglandins are key to establish the priming response, and lipoxins maintain a broad general state of enhanced immune surveillance that is not pathogen-specific. This begs the question of whether eicosanoids may also be important mediators of trained immunity in humans.

## Acknowledgments

The authors gratefully acknowledge Asher Kantor and Mark Johnson for editorial assistance.

## Ethics Statement

Animal studies were done according to the NIH animal study protocol (ASP) approved by the NIH Animal Care and User Committee (ACUC), with approval ID ASP-LMVR5. Public Health Service Animal Welfare Assurance #A4149-01 guidelines were followed according to the National Institutes of Health Animal (NIH) Office of Animal Care and Use (OACU).

