## Supplementary Methods and Figures for "“Proliferation of DBLOX Peroxidase-Expressing Oenocytes Maintains Innate Immune Memory in Primed Mosquitoes”"

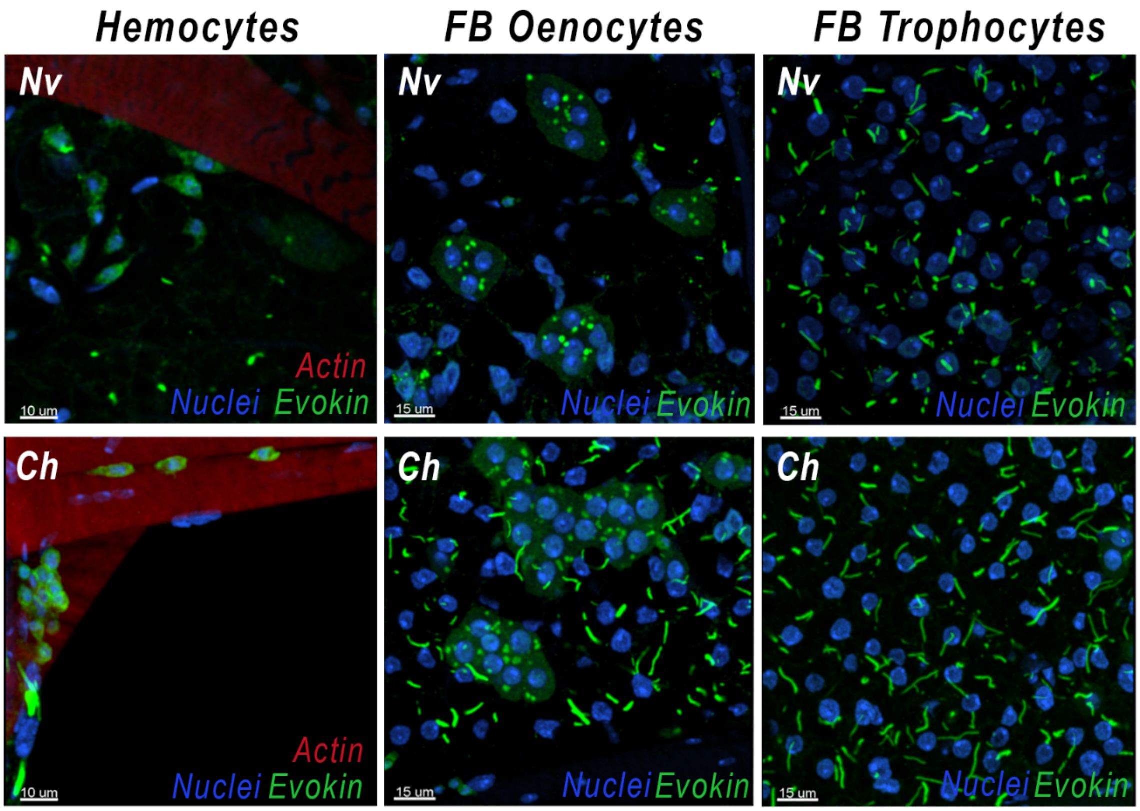


**Fig. S1 – Evokin staining in blood-fed naïve (Nv) and *P. berghei* challenged (Ch) mosquitoes.** Sessile body wall attached hemocytes, oenocytes, and trophocytes 7 days post infection. Nuclei (blue), Evokin (green), actin (red). Scale bars: 10 and 15 µm.

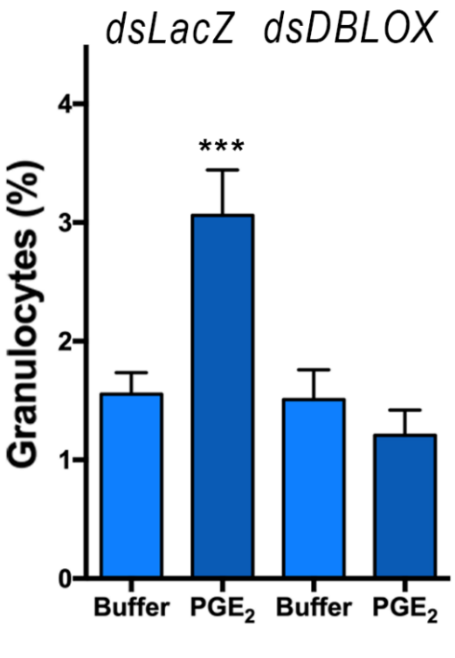


**Fig. S2 – Effect of silencing DBLOX on PGE2-injection induced granulocyte priming.** Mean ± SEM are plotted and groups were compared using Student's t test (***p<0.001).

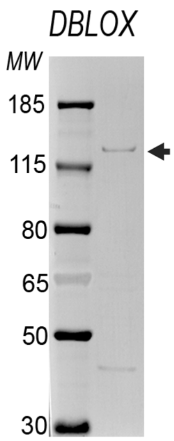


**Fig. S3. DBLOX protein expression in mosquito body wall analyzed by Western Blot.** Mosquito body walls were collected from *P. berghei* infected females 7 days post-feeding. Homogenates were analyzed under denaturing and reducing conditions. DBLOX protein was detected with a rabbit anti-DBLOX antibodies. The arrow indicates the main band with a molecular weight corresponding to that predicted for DBLOX (159 kDa).

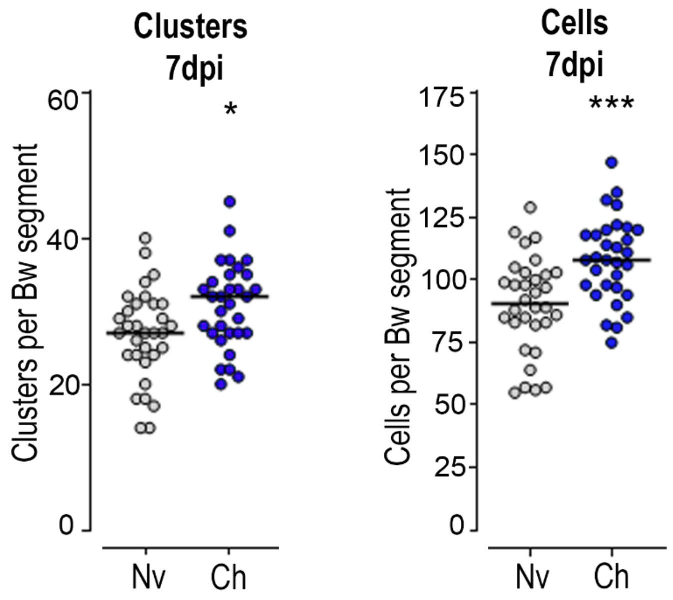


**Fig. S4 – Proliferation of DBLOX-expressing oenocytes in response to *P.berghei*.** Number of DBLOX-expressing oenocyte clusters per body wall segment (left), and number of DBLOX-expressing oenocytes per body wall segment (right) 7 days post infection.. Lines indicate medians that were compared using the Mann-Whitney U test. (*p<0.05, ***p<0.001).

| Gene | Forward | Reverse |
|---|---|---|
| **qRT-PCR** | | |
| HPX1 | GTTGGCGGAGAAGATAGCGA | GCCACATTCACTGCATCGAC |
| HPX2 | CCGCTTCTACAACACGATGA | CGACCAGATGGGCAAGTAT |
| HPX3 | CCTCGAACCAGACGGAGATG | TCAGCGATATTTTCGGCGGT |
| HPX4 | GAACAGTGCGATGCGTCAAA | TTTCGTCAGCTGGTTCTCGT |
| HPX5 | CGCCTTCGACTACAACCTCA | GTCTTCCAGTCACCCTTCGG |
| HPX6 | GGTCGGGGCAGAAATGCAG | GCGTTGTGTACGATTCGACC |
| HPX7 | ACTGTACCCTCGAAGCTGGA | GCGTGGGATTGAGAAAGGTA |
| HPX8 | AGTCACGTACGCATCGTCAG | ACATCACTCGCAGGACACAG |
| HPX10 | AGACGCTGAATCCAACCTGG | AAATTGCGACCCAGAAACGC |
| HPX11 | GTTGCCCAACCCTCGAAAAC | GCGTTAGGGCCATATCGTGA |
| HPX12 | ACAACCGTCCAGTCTGTGTG | ATGCGCAAGTAGCAGACGTA |
| HPX13 | GCTACGTGGACCAAGAGACA | GGATGTCGTTCAGCTGTGGT |
| HPX14 | TTTCAATGGCACTGTTCGGC | CGGTGGAATTCGTTTACCGC |
| HPX15 | GTTCAACAACCCTTCGGTGC | TGCCGGAAGAGGAAATGCTT |
| HPX16 | GATGCCGTTCATGCAATCCC | ATCGACCTGTTTGGCCGATT |
| DBLOX | GCTGAGTTCGGAAGACGAC | CAATTTGTGGCATGGAGTTG |
| DUOX | GCCCGCAGGAAGTTCGTCAAAAA | TCGCATCATCTCGCTCAGTTCTCC |
| Evokin | CAGCCGAAAGTGAACAAACA | ATGGTGGCCATTGTATACGG |
| S7 | AGAACCAGCAGACCACCATC | GCTGCAAACTTCGGCTATTC |
| **dsRNA** | | |
| AgTip60 | taatacgactcactatagggCACCGGATCAAACCGTGGTA | taatacgactcactatagggTATGCACGCCACGTTGTAGT |
| AGAP000029 | taatacgactcactatagggAACAAGTCGAAGGCAGCTCA | taatacgactcactatagggGTGGCCCACCTTCTCCTTAC |
| AGAP004434 | taatacgactcactatagggGCCAACACTCTGGAACGGTA | taatacgactcactatagggAAACAACGACGGTCATCGGA |
| AGAP003264 | taatacgactcactatagggCCAAAAACTGGGCATTGCGA | taatacgactcactatagggGCACCTGGCCATAGATCTCC |
| AGAP012229 | taatacgactcactatagggAAAGGAGCACGAAGCGATCA | taatacgactcactatagggACAGCTTCGCCATCAGACA |
| AGAP002735 | taatacgactcactatagggGACCAGTGCGAAGATAGCGA | taatacgactcactatagggTTGCTGCGGCTTGTTTTACC |
| AGAP002231 | taatacgactcactatagggCACAGCAAAAGTCATCGCCC | taatacgactcactatagggCCGATTAAGCATTGCGCTGT |
| AGAP009676 | taatacgactcactatagggGGTCGATTCCGGTGTACGAA | taatacgactcactatagggAATTTGGCCAGCAAGCACAG |
| AGAP008300 | taatacgactcactatagggCGGGACGGTACAAGAGCTAC | taatacgactcactatagggCCCGATATAACCGCCAGCTT |
| AGAP002568 | taatacgactcactatagggGGCCAAGCTTGCCTACTACA | taatacgactcactatagggATGGCTGTCTCCATTCGCAA |
| HPX2 | taatacgactcactatagggACGACGACGGTGTGTACAAG | taatacgactcactatagggATACTCGGCCGAATCGAAC |
| DBLOX | taatacgactcactatagggCGGACTGGTGGACGAGATTTC | taatacgactcactatagggCGGACTGGTGGACGAGATTTC |
| **Cloning** | | |
| **DBLOX** Domain 2 | AAGGAGATATACATATGAACCAGTTCATT ACCGGGTCGATCT | GCCGGATCTGCTCGAGTCAATGGTGGTGATGATGATGGTGAC ACTGGGGTCCCTCGTTGGT |
| Evokin (INFEVO17 | AAGGAGATATACATATGCAGGTTCCGGG TTTC | GCCGGATCTGCTCGAGTTAATGATGATGATGATGATGGCAGTT TTTCTGGTCG |

**Supplemental Table 1: List of primer used for qRT-PCR, dsRNA synthesis and cloning.**

### Materials and Methods

#### Animals, Priming and Granulocyte Counting

*A. gambiae* (G3 strain) mosquitoes were reared at 28°C, 80% humidity, 12h light/dark cycle. Mosquitoes were fed with 10% Karo syrup solution *ad libitum*. For *Plasmodium* infection-induced priming, the *P. berghe*i Anka GFP strain was used. Mice infections were performed through serial passage in 4-5 week-old female BALB/c mice. A drop of blood was taken from the mouse tail 48-72h after passage, and parasitemia was evaluated on Giemsa-stained, methanol-fixed blood smear slides. For mosquito infections, mice carrying 3-5% parasitemia infections were chosen. After infection, mosquitoes were kept at 19°C, 80% humidity, 12h light/dark cycle until the day of sample processing. Animal studies were done according to the NIH animal study protocol (ASP) approved by the NIH Animal Care and User Committee (ACUC), with approval ID ASP-LMVR5. Public Health Service Animal Welfare Assurance #A4149-01 guidelines were followed according to the National Institutes of Health Animal (NIH) Office of Animal Care and Use (OACU).

Alternatively, mosquitoes were primed by injection of cell-free hemolymph transfer from challenged mosquitoes as previously described. Hemolymph was collected from pools of 10 females perfused with 10µl of a modified anticoagulant buffer (95% Schneider's medium and 5% citrate buffer). The pool of hemolymph was centrifuged at 9,300g for 10 min at 4°C and the cell-free supernatant was transferred and stored at -80°C. Cell-free hemolymph challenge was done by injecting 138nL of cell-free hemolymph into a 3 days-old sugar-fed naive mosquito. For $PGE_2$ priming, an aliquot of ethanol-diluted $PGE_2$ was dried in nitrogen gas stream, resuspended in anticoagulant buffer and $PGE_2$ was injected into 3 days-old sugar-fed naive mosquitoes.

For granulocyte counting, hemolymph was collected from females perfused with 10µl of hemocyte counting buffer (10% fetal bovine serum, 30% citrate buffer in Schneider's medium). Collected samples were applied to sterile hemocytometer slides and granulocytes were counted in a light microscope as the relative number of cells against total hemocyte numbers.

#### Sua Cell Culture and Priming

Sua cells were grown on Sua Cell Media (10% fetal bovine serum, 1,000 I.U./mL Penicillin, 1 mg/mL Streptomycin on Schneider's media) at 28°C and passed at 80-90% confluence. For bacteria challenge,

10mg/ml *Escherichia coli* acetone powder was added to the cell media and cells were incubated for 20 hours. Following, cells were extensively washed with cell media and incubated overnight. On the next day, media was removed and cells were processed.

### Gene Expression Analysis

For gene expression analysis, midgut and abdominal fat body specimens from 15-20 females per group were dissected. RNA samples from dissected midguts were obtained using the RNeasy RNA extraction kit (Qiagen) following standard protocol. Hemolymph samples were prepared by perfusing mosquitoes with 5 µl of a modified anticoagulant buffer (95% Schneider's medium and 5% citrate buffer) into TRIzol (Invitrogen). Fat body RNA and Sua cell samples were prepared by standard TRIzol (Invitrogen) protocol. The RNA concentration was measured using NanoDrop (Thermo Scientific) and cDNA samples were prepared with the QuantiTect reverse transcription kit with DNA wipeout (Qiagen) following standard protocol. Relative gene expression was analyzed by qRT-PCR using the DyNamo SYBR green qPCR kit (Thermo Scientific) and a CFX96 Real-Time PCR Detection System (BioRad). Data analysis was done using the ΔΔCt method.

### Sua Cell Protein Fractionation

For Sua cell protein fractionation, the cell media was removed and cold 0.2 mM PMSF, PBS was added. Cells were scraped and centrifuged (5 min, 5,000 rpm, 4$^{o}$C). The pellet was resuspended in protease inhibitor supplemented cold Buffer A (1.5 mM MgCl2, 10 mM KCl, 0.5 mM DTT, 0.2 mM PMSF, 10 mM Hepes pH 7.9) and incubated on ice for 10 min. Following, cells were vortexed and centrifuged (5 min, 5,000 rpm, 4$^{o}$C). The supernatant (cytoplasmic fraction) was stored. The pellet was resuspended in protease inhibitor supplemented cold Buffer C (25% glycerol, 420 mM NaCl, 1.5 mM $MgCl_2$, 0.2 mM EDTA, 0.5 mM DTT, 0.2 mM PMSF, 20 mM Hepes pH 7.9), incubated on ice for 20 min and centrifuged (10 min, 20,000g, 4$^{o}$C). The supernatant (soluble nuclear fraction) was stored and the pellet (membrane fraction) was resuspended in Solubilization Buffer (150 mM NaCl, 5mM EDTA, 2% SDS, 1 mM Tris-HCl pH 7.5) and stored at -70$^{o}$C.

### Recombinant Double Peroxidase Protein Expression and Immunization

A region of the second domain of the full-length *A. gambiae* Double Peroxidase (AGAP008350/DBLOX) coding sequence was amplified and subcloned into a pET17B plasmid containing a hexahistidine tag at the C terminus. The Double Peroxidase Domain 2 construct was expressed in One Shot™ BL21(DE3)pLysE T7 Competent *E. coli* (ThermoFisher Scientific). *E. coli* transformed with the Double Peroxidase Domain 2 construct were grown at 37°C with vigorous shaking until reaching an optical density of 0.4-0.6, at which point 1 mM isopropyl β-D-thiogalactopyranoside (IPTG) (Sigma) was added to induce expression for 4 hours. Pelleted *E. coli* were sonicated in a solution containing 8 M urea in HEPES-buffered saline (20 mM HEPES pH 7.5, 150 mM NaCl) to solubilize inclusion bodies, then purified in a column using HisPur™ Ni-NTA Resin (ThermoFisher Scientific) for affinity chromatography. Purified recombinant Double Peroxidase protein was run in an SDS-PAGE, excised from the gel, and immunized in rabbit.

### Recombinant Evokin Protein Expression and Immunization

The mature full-length AGAP009281-PA coding sequence was codon-optimized for *Escherichia coli* cell expression and synthesized by BioBasic (Amherst, NYJ). The AGAP009281-PA construct was built by PCR amplification and was sub-cloned by In-Fusion (Clontech) into a pET17 vector (EMD Millipore) between Nde I and Xho I sites, with primers INFEVO17 (Supplemental File). The construct was expressed in BL21 Escherichia coli (Invitrogen). HEPES-buffered saline (20 mM HEPES pH 7.5, 150 mM NaCl) 8M urea solution containing inclusion bodies of cultures induced for 3 h at 37◦C with 1 mM isopropyl β-D-thiogalactopyranoside (Sigma) were purified by nickel affinity chromatography and stored after being dialyzed in HEPES-buffered saline.

#### Antibody generation

All animal procedures were performed according to protocols approved by the NIAID and NIH Animal Care and Use Committee. Female Balb/c mice, 5–8-week-old naïve, were purchased from Charles River (Germantown, MD) and maintained at a facility at the NIH. Groups (n =5) of female BALB/c mice were immunized subcutaneously in their ears with a 20 µL solution that contained 5 µg of HEPES-buffered saline-dialyzed recombinant AGAP009281-PA protein emulsified in Magic mouse adjuvant (Creative Diagnostics # CDN-A001). The immunized mice were boosted twice at 2-week intervals with the same quantity of protein. Blood was collected on day 0 (Pre-immune sera) and 2 weeks after each subsequent immunization for analysis of antibodies titer or terminal bleeding.

### Western Blot

Homogenate samples were boiled in NuPAGE™ LDS Sample Buffer (ThermoFisher) and separated using a 12% Bis-Tris gel. The proteins were transferred to a nitrocellulose (Evokin) or a PVDF membrane (DBLOX), blocked for 4 hours in blocking buffer (5% non-fat dry milk, 0.1% Tween 20, TBS). Proteins were incubated with primary antibodies diluted in blocking buffer at the following concentrations: anti-Evokin - 1:1000; anti-DBLOX - 1:2000. The membranes were subsequently washed three times in TBST and incubated with goat anti-mouse or anti-rabbit secondary antibodies conjugated to alkaline phosphatase at a 1:5000 dilution. The membranes were developed using Western Blue® Stabilized Substrate for Alkaline Phosphatase (Promega).

To remove unspecific antibodies that could target for *E. coli* proteins, Double Peroxidase serum was diluted and preabsorbed twice for 30 mins with *E. coli* acetone powder immediately before it was used. Briefly, One Shot™ BL21(DE3)pLysE T7 Competent E. coli (ThermoFisher) was grown overnight at 37$^{o}$C, pelleted, and resuspended on 0.9% NaCl. The *E. coli* were lysed and dehydrated using acetone, pelleted, and crushed into a fine powder and washed tree times with PBS. For pre-absorption, 1% w/v of the powder pellet was used.

### Immunohistochemistry

Mosquitoes were individually injected with 207 nL of 16% paraformaldehyde and left for 20 seconds. Then in 1X PBS, the mosquito abdomen was separated from the head and thorax, and the midgut, ovaries, and Malpighian tubes were removed. The abdomens were cut longitudinally through the dorsal region resulting in a flat, open tissue with an exposed fat body. Opened abdomens were immediately placed into 4% paraformaldehyde and were left to fix for 30 mins. The tissues were washed twice with 1X PBS and then delipidated in ice-cold ethanol in a stepwise manner starting with 20% up to 80% in 20% intervals, then back down. Tissues were washed twice with 1X PBS and then placed into 4% paraformaldehyde and left for 1 hour with gentle shaking. Fixed tissues were washed twice with 1X PBS, washed three times with PBST (0.1% Triton) for 5 minutes each. Tissues were blocked in blocking buffer (0.1% Triton, 2% BSA, 0.1% gelatin, in PBS) for 2 hours at room temperature and incubated overnight at 4$^{o}$C with the Rabbit DBLOX primary antibody diluted 1:1500 in blocking buffer and preabsorbed with *E. coli* acetone powder twice, as previously described. Tissues were washed with blocking buffer and then blocked in blocking buffer with 20% goat serum for 1 hour and then incubated for 2 hours room temperature with a goat anti-rabbit Alexa Fluor 594 secondary antibody (Fisher Scientific) diluted 1:1000 in blocking buffer with 20% goat serum. Tissues were washed with blocking buffer and then PBST and labeled with Phalloidin 488 (1:40) and Hoechst (1:10000) for 30 mins and then incubated in PBS + 10% glycerol for 20 mins before mounting on slides in Prolong Gold mounting media (Molecular Probes).

#### dsRNA Synthesis and Gene Silencing

To prepare dsRNA for gene silencing, a cDNA template of challenged mosquitoes was first amplified by PCR using primers with flanking T7 promoter recognition sequences (Supplementary Table 1). Following amplification, PCR products were purified using the QIAquick PCR purification kit (Qiagen) and double stranded RNA was synthetized using the MEGAscript RNAi kit (Ambion) according to manufacturer's instructions. For gene silencing, 69-128 nL of 3 µg/µl dsRNA was injected in 2-3 day-old naive females.
